# Predictive markers for Parkinson’s disease using deep neural nets on neuromelanin sensitive MRI

**DOI:** 10.1101/523100

**Authors:** Sumeet Shinde, Shweta Prasad, Yash Saboo, Rishabh Kaushick, Jitender Saini, Pramod Kumar Pal, Madhura Ingalhalikar

## Abstract

Neuromelanin sensitive magnetic resonance imaging (NMS-MRI) has been crucial in identifying abnormalities in the substantia nigra pars compacta (SNc) in Parkinson’s disease (PD). PD is characterized by loss of dopaminergic neurons in SNc and current techniques employ estimation of contrast ratios of the SNc visualized on NMS-MRI to discern PD patients from the healthy controls. However, the extraction of these features is time-consuming and laborious and moreover provides lower prediction accuracies. Furthermore, these do not account for patterns of subtle changes in PD in the SNc. To mitigate this, our work establishes a computer-based analysis technique based on convolutional neural networks (CNNs) to create prognostic and diagnostic biomarkers of PD from NMS-MRI. The technique not only performs with a superior cross-validation accuracy (83.7%) as well as testing accuracy (80%) as compared to contrast ratio-based classification (52.7% cross-validation and 56.5% testing accuracy) and radiomics based classifier (81.1% cross-validation and 60.3% testing accuracy); but also locates the most discriminative regions on the neuromelanin contrast images. These discriminative activations demonstrate that the left SNc plays a key role in the classification in comparison to the right SNc, and are in agreement with the concept of asymmetry in PD. Overall, the proposed technique has the potential to support radiological diagnosis of PD while facilitating deeper understanding into the abnormalities in SNc.

## Introduction

Parkinson’s disease (PD) is progressive, neurodegenerative disorder characterized by loss of dopaminergic neurons in the substantia nigra pars compacta (SNc) (Mann & Yates, 1983). Neuromelanin which is a by-product of dopamine synthesis is a neuronal pigment which can be visualized in the SNc. Depigmentation of the SNc secondary to loss of dopaminergic neurons is a conspicuous feature of PD, which although well visualized neuropathologically is poorly replicated by neuroimaging(Hutchinson & Raff, 2000). The role of conventional neuroimaging in the diagnosis of PD has been therefore limited, despite several methods to study the SNc since these techniques have been unable to directly visualize SNc. The introduction of the “NM-sensitive MRI”, a 3T T1 weighted high-resolution fast spin-echo neuromelanin sensitive sequence by Sasaki et al revolutionized the technique of visualizing the SNc (Sasaki et al., 2006). Following this, several studies have demonstrated the utility of this sequence to differentiate between patients with PD and controls using manually extracted features of the SNc that include the contrast ratio, area and volumes(Matsuura et al., 2013; Ogisu et al., 2013; Ohtsuka et al., 2013; Prasad, Stezin, et al., 2018). Between these features the contrast ratios are well-accepted as they have demonstrated the highest discriminative power. However, these techniques have limited clinical utility and reproducibility owing to the time-consuming nature of computing these features, with a high scope for operator errors. Moreover, these features require SNc borders to be well defined which are difficult in patients with PD due to advanced loss of neuromelanin containing neurons that may substantially decrease the contrast on the scan and may bias the analysis by excluding highly impacted areas of the SNc, leading to an over-estimate in contrast ratios(Sulzer et al., 2018).

At present the diagnosis of PD is highly dependent on clinical features and although dopamine transporter positron emitted topography is useful, it is not cost-effective and cannot be routinely employed. Hence, it is crucial to employ other neuro-imaging techniques to aid in the early or differential diagnosis of PD. Recent advances in the areas of machine learning and data-driven analysis have demonstrated the utility of different brain imaging modalities for automated diagnosis of PD. These studies have utilized a host of techniques that include supervised predictive models such as support vector machines (SVMs) (Abos et al., 2017; Amoroso, La Rocca, Monaco, Bellotti, & Tangaro, 2018; Cherubini, Morelli, et al., 2014; Cherubini, Nistico, et al., 2014; Huppertz et al., 2016; Rana et al., 2015; Salvatore et al., 2014) as well as unsupervised models such as self-organizing maps (Peran et al., 2018; Singh & Samavedham, 2015) on data acquired from morphological T1 weighted MRI, functional MRI, diffusion tensor imaging, SPECT, etc. (Adeli et al., 2016; Ariz et al., 2018) and have reported high but variable accuracies. Table 1 provides a brief review of recent articles that have used machine learning and statistical learning techniques to predict PD from MRI modalities.

**Table 1:** Brief review of methods employed by recent studies that have used machine learning and statistical learning techniques to predict PD from MRI modalities

| Author, year | Number of subjects | Methods employed |
| --- | --- | --- |
| Salvatore et al, 2013 | PD (n=28)<br>PSP (n=28)<br>HC (n=28) | VBM<br>Principal component analysis<br>SVM |
| Cherubini et al, 2014 | Tremor dominant PD (n=15)<br>ET with rest tremor (n=15) | VBM, DTI<br>SVM |
| Cherubini et al, 2014 | PD (n=57)<br>PSP (n=21) | VBM, DTI<br>SVM |
| Rana et al, 2015 | PD (n=30)<br>HC (n=30) | Region of interest based<br>SVM |
| Singh and Samavedham 2015 | PPMI cohort<br>PD (n=518)<br>SWEDD (n=68)<br>HC (n=245) | Self-organizing maps<br>SVM |
| Huppertz et al, 2016 | PD (n=204)<br>PSP-RS (n=106)<br>MSA-C (n=21)<br>MSA-P (n=60) | Volumetry<br>SVM |
| Adeli et al, 2016 | PPMI cohort<br>PD (n=374)<br>HC (n=169) | Joint feature-sample selection |
| Abos et al, 2017 | PD (n=27)<br>HC (n=38) | Functional connectome<br>SVM |
| Peran et al, 2018 | PD (n=26)<br>MSA-P (n=16)<br>MSA-C (n=13)<br>HC (n=26) | VBM, T2* relaxometry, DTI<br>Self-organizing maps |
| Amoroso et al, 2018 | PPMI cohort<br>PD (n=374)<br>HC (n=169) | Connectivity measures<br>SVM |
| Ariz et al, 2018 | PD (n=40)<br>HC (n=39) | NM-MRI based atlas of Substantia nigra |

DTI: Diffusion tensor imaging; ET: Essential tremor; HC: Healthy controls; MSA-C: Multiple system atrophy with predominant cerebellar features; MSA-P: Multiple system atrophy with predominant parkinsonian features; NM-MRI: Neuromelanin sensitive magnetic resonance imaging; PD: Parkinson's disease; PPMI: Parkinson's Progression Markers Initiative; PSP: Progressive supranuclear palsy; PSP-RS: Progressive supranuclear palsy-Richardson syndrome; SVM: Support vector machine; SWEDD: Scans without evidence of dopaminergic deficit; VBM: Voxel based morphometry

Considering the above-mentioned utility of the NMS-MRI in the differentiation of PD from healthy controls, and the relative ease of acquiring this sequence, we endeavor to utilize this sequence in an automated classification framework for PD prognosis and diagnosis. The main objective of the present study therefore is to create markers for PD using state-of-art deep convolution neural network (CNN) on NMS-MRI and compare it to classifiers based on (1) contrast ratios with machine learning (CR-ML) and (2) radiomics with machine learning (RA-ML) using regions of interest of the SNc. Our CNN based classifier is fully automated and is free of defining SNc borders as it uses a larger region of interest. Moreover, it employs the technique of discriminative localization to compute class activation maps (CAMs) that facilitate deeper understanding into the most important regions that participate in the classification.

## Methodology

### Subject recruitment and clinical evaluation

Forty-five patients with PD and 35 healthy controls (HCs) were recruited from the general outpatient clinic and movement disorder services at the Department of Neurology, National Institute of Mental Health and Neurosciences (NIMHANS), Bangalore, India. The diagnosis of idiopathic PD was based on the UK Parkinson’s Disease Society Brain Bank criteria (Hughes, Daniel, Blankson, & Lees, 1993) and confirmed by a trained movement disorder specialist (author PKP). Patients included in this study have been part of other studies from this group (Prasad, Saini, Yadav, & Pal, 2018; Prasad, Stezin, et al., 2018) and all patients and controls provided informed consent prior to recruitment in the original projects.

Demographic and clinical details such as gender, age at presentation, age at onset of motor symptoms, disease duration, Unified Parkinson’s Disease Rating Scale (UPDRS-III) OFF-state scores and Hoehn and Yahr stages were recorded. Age and gender matched HCs with no family history of parkinsonism or other movement disorder were recruited.

### Imaging

MR images were acquired on 3T Philips Achieva scanner with a 32-channel head coil at NIMHANS, Bangalore, India. High resolution 3D neuromelanin contrast sensitive sequence i.e. the spectral pre-saturation with inversion recovery (SPIR) sequence was acquired using TR/TE: 26/2.2ms, flip angle: 20°; reconstructed matrix size: 512 x 512; field of view: 180×180×50mm; voxel size: 0.9×0.9×1mm; number of slices: 50. with an acquisition time 4 min 12.9 s. The MR images were retrieved from the archive and screened for gross cortical structural abnormalities by an experienced neuroradiologist (author JS), after which the images were set perpendicular to the fourth ventricle floor with coverage between the posterior commissure and inferior border of the pons.

### Convolutional Neural Nets with Discriminative Localization (CNN-DL)

CNNs are a modern adaption of the traditional artificial neural network architecture where millions of 2D convolutional filter parameters are computed from multiple levels of granularity and transformed into the desired output by end-to-end optimization(Krizhevsky, Sutskever, & Hinton, 2012). The CNN can therefore be considered as an automated feature extraction tool with a classifier in the final stages.

In our case, we employed a boxed region around the brain-stem on the axial slices of the NMS-MRI as input to the 2D CNN (Figure 1). The CNN architecture was derived from ResNet50 design (He, Zhang, Ren, & Sun, 2016), which is a highly robust design and has been shown to work with superior performance on medical image classification tasks such as in tumor classification(Chang et al., 2018; Korfiatis et al., 2017). ResNets, unlike other CNN architectures, consistof short-cut connectionsthat facilitate learning of residuals and ensure that the subsequent layers have the necessary information to learn additional features (Ferreira et al., 2006). Our design (as shown in Figure 1) consists of16 residual blocks, each of which contains 3 convolutional layers and an identity connection. A max-pooling layer, that downsamples the image, is applied before the first block and after 3,4,6 and 3 residual blocks as shown in Figure 1. The convolutional layers contain 3*3 kernels that capture the patterns and features of varying granularity depending upon the depth of the layer. The last convolutional layer acquires the most abstract markers that is given as input to the global average pooling (GAP) layer which averages across the output of the convolutional layer for both the classes (PD and controls in our case) resulting in 2 representations, one for each class using a softmax activation.

**Figure 1:**
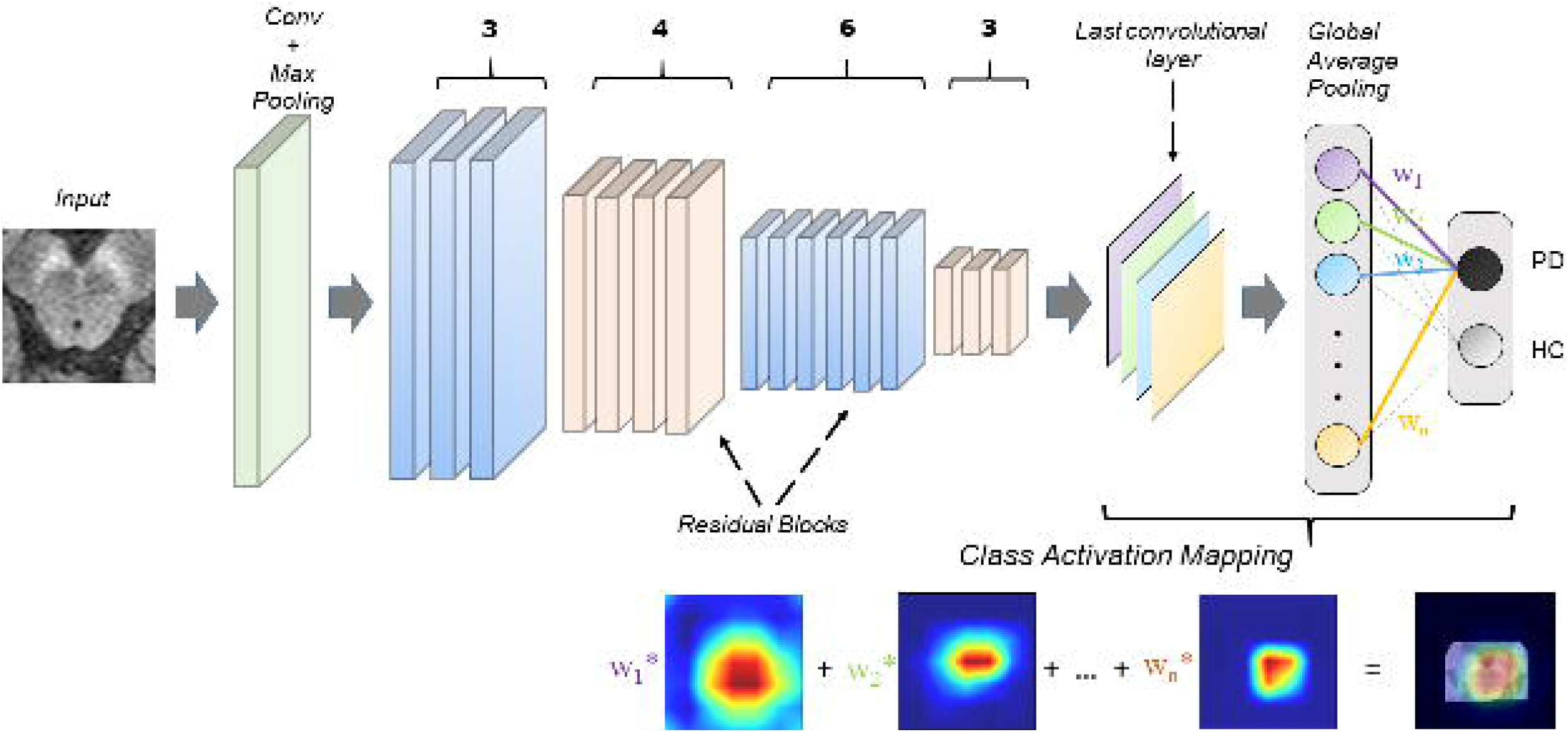
The figure displays a schematic diagram of the CNN architecture. ResNet50 architecture was employed with 16 blocks (50 layers in total). The class activation maps (CAMs) were computed using global average pooling as shown in the figure.

Throughout the CNN architecture, we employ rectified linear unit (ReLU) activations and with a learning rate of 0.0001 which decays based on the number of iterations. The weights of each of layers are updated to train the complete CNN model by minimizing the categorical cross entropy loss function as given in equation 1 where *y_n_* is target output probability, 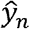 is predicted output probability, *N* is number of testing samples and *J*(*w*) is categorical cross-entropy loss, and is achieved by using the Adam optimizer. Finally, to reduce the susceptibility to over-fitting, as a standard practice, data augmentation was performed which increased the dataset by several folds.

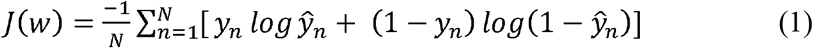

To obtain the discriminative activations, we forward propagate the input image and acquire the weights (*w_1_, w_2_, w_3_… w_n_*) for the output layer, as given in Zhou et al. (Zhou, Khosla, Lapedriza, Oliva, & Torralba, 2016). The feature maps (*f_1_, f_2_, f_3_… f_n_*) of the last convolutional layer are also obtained and are upsampled to match the resolution of the original input image. To create the class activation map, the weights for the predicted class are multiplied with the corresponding feature maps and then added together. The resulting map can therefore demonstrate the most discerning regions in the image for each subject. Finally, to assess any contra-lateral deficits in PD, as have been shown in earlier studies, for each correctly classified subject, the mean activations were computed separately on the left and right, by dividing the input image into two parts.

### Contrast Ratio computation

To compute the contrast ratio, technique described in Prasad et al.(Prasad, Stezin, et al., 2018) was employed. A section of the midbrain at the midpoint of the mamillary bodies was selected for placement of ROIs. Signal intensity (SI) was measured by placing 10mm^2^ circular ROIs over the lateral part of bilateral SNc, and a normative SI (SI_N_) was obtained by placing a ROI anterior to the cerebral aqueduct. Contrast ratios (CRs) of the SNc were calculated based on the methodology described by Ohtsuka et al. (Ohtsuka et al., 2013). The following equation was used for the calculation of the CRs:

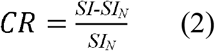

where SI is the value of the SI of either right or left SNc and SI_N_ is the SI of the region anterior to the cerebral aqueduct.CR for right and left SNc was computed separately. The left and right contrast ratios were employed in a random forest (Breiman, 2001) with XGBoost classification [https://xgboost.readthedocs.io/en/latest/], a gradient boosting framework, for comparison with our CNN model. The complete schematic procedure is shown in the supplementary material-figure 1.

### Radiomics

Radiomics involves extraction of a large number of quantitative measures such as textures, intensity features, gray-level co-occurrences, gray-level run lengths, statistical measures and energy features (Gillies, Kinahan, & Hricak, 2015). For our analysis, we employed the PyRadiomics (van Griethuysen et al., 2017) toolbox and computed 1470 number of features (details of which are provided in supplementary material -table 1) from the 3D SNc right and left region of interests. Bilateral SNc ROIs were created by manual segmentation of the NMS-MRI. An expert neurology trainee (author-SP) who was blinded to the groups, delineated the right and left SNc on the axial slices and created a 3D binary mask. The radiomic measures were normalized using min-max normalization and were used as features in a random forest classifier (Breiman, 2001) with XGBoost for classification between PD and healthy controls. This was performed as a comparison with our proposed CNN model. Ten top ranked (most discriminative) radiomic features were computed using the average information gain for each feature over all the decision trees (f-score), to facilitate deeper understanding of the patterns of abnormality from the NMS-MRI images in PD. Supplementary material -figure 1 shows the schematic diagram for the procedure employed.

### Training and Testing of the models

The dataset was randomly divided into training and testing while retaining the PD:HC ratio in both the classes. We used, 25 healthy controls and 30 PD subjects for training and crossvalidating (5-fold validation) the models (our proposed CNN-DL and the comparative CR-ML and RA-ML) and the remaining 10 HCs and 15 PD patients (25 subjects) for testing the models. The receiver operation characteristics (ROC), accuracy, F1-score, precision and recall (Powers, 2011) were computed on the cross-validation and test dataset and reported.

## Results

Table 2 provides complete demographic information of the dataset under consideration. There were no significant differences observed between the age and gender of the patient and control group. The UPDRS (OFF) scores, duration of illness and age at onset are reported. With the exception of a single patient with PD, all other subjects were right-handed. Our CNN-DL classifier performed with a cross-validation accuracy of 83.7% (AU-ROC = 0.90) and test accuracy of 80% (AU-ROC = 0.91), while the CR-ML performed with a cross-validation accuracy of 52.7% (AU-ROC = 0.47) and a test accuracy of 56.5 % (0.54 AU-ROC), while RA-ML performed with a cross-validation accuracy of 81.1% (AU-ROC = 0.89) and 60.3% test accuracy (0.54 AU-ROC). Figure 2 shows the ROC curves for the CNN-DL in comparison with the other two methods. The testing accuracy, F1-score, precision and recall are reported in Table-3. The top-most radiomics based features included the run length features, non-uniformity, surface-volume ratio, grey level emphasis as shown in figure 3.

**Table 2:** Demographic details of patients with Parkinson’s disease and healthy controls

|  | Parkinson’s disease<br>(n=45) | Healthy controls<br>(n=35) |
| --- | --- | --- |
| Gender (Male: Female) | 32:13 | 22:13 |
| Age | 58.00±8.70 | 55.00±5.40 |
| Age at onset | 51.90±8.64 | - |
| Duration of illness | 6.26±4.06 | - |
| UPDRS III (OFF) scores | 36.58±13.66 | - |

UPDRS: Unified Parkinson's disease rating scale

**Figure 2:**
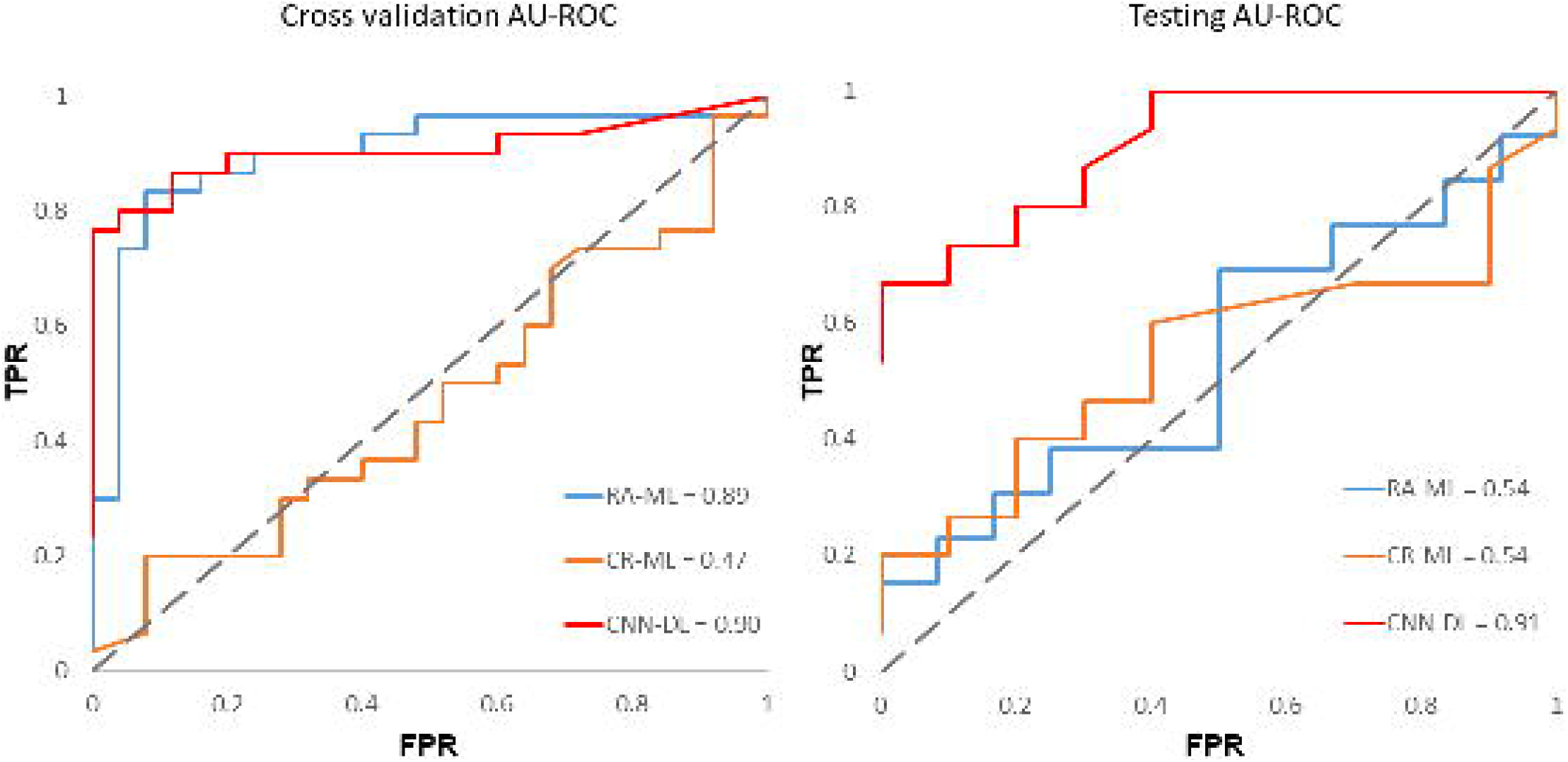
Receiver operating characteristics for all the three methods employed (a) crossvalidation (b) testing.

**Table 3:** Performance of convolution neural networks compared to contrast ratios with machine learning (CR-ML) and radiomics with machine learning (RA-ML)

|  | CR-ML | RA-ML | CNN-DL |
| --- | --- | --- | --- |
| Cross Validation results |  |  |  |
| Precision | 0.56 | 0.92 | 0.78 |
| Recall | 0.73 | 0.86 | 0.88 |
| F1-score | 0.628 | 0.830 | 0.836 |
| Accuracy | 52.7% | 81.1% | 83.7% |
| AU-ROC | 0.47 | 0.89 | 0.90 |
| Testing results |  |  |  |
| Precision | 0.72 | 0.64 | 0.85 |
| Recall | 0.53 | 0.69 | 0.60 |
| F1-score | 0.592 | 0.642 | 0.705 |
| Accuracy | 56.5% | 60.3% | 80.0% |
| AU-ROC | 0.54 | 0.54 | 0.91 |

**Figure 3:**
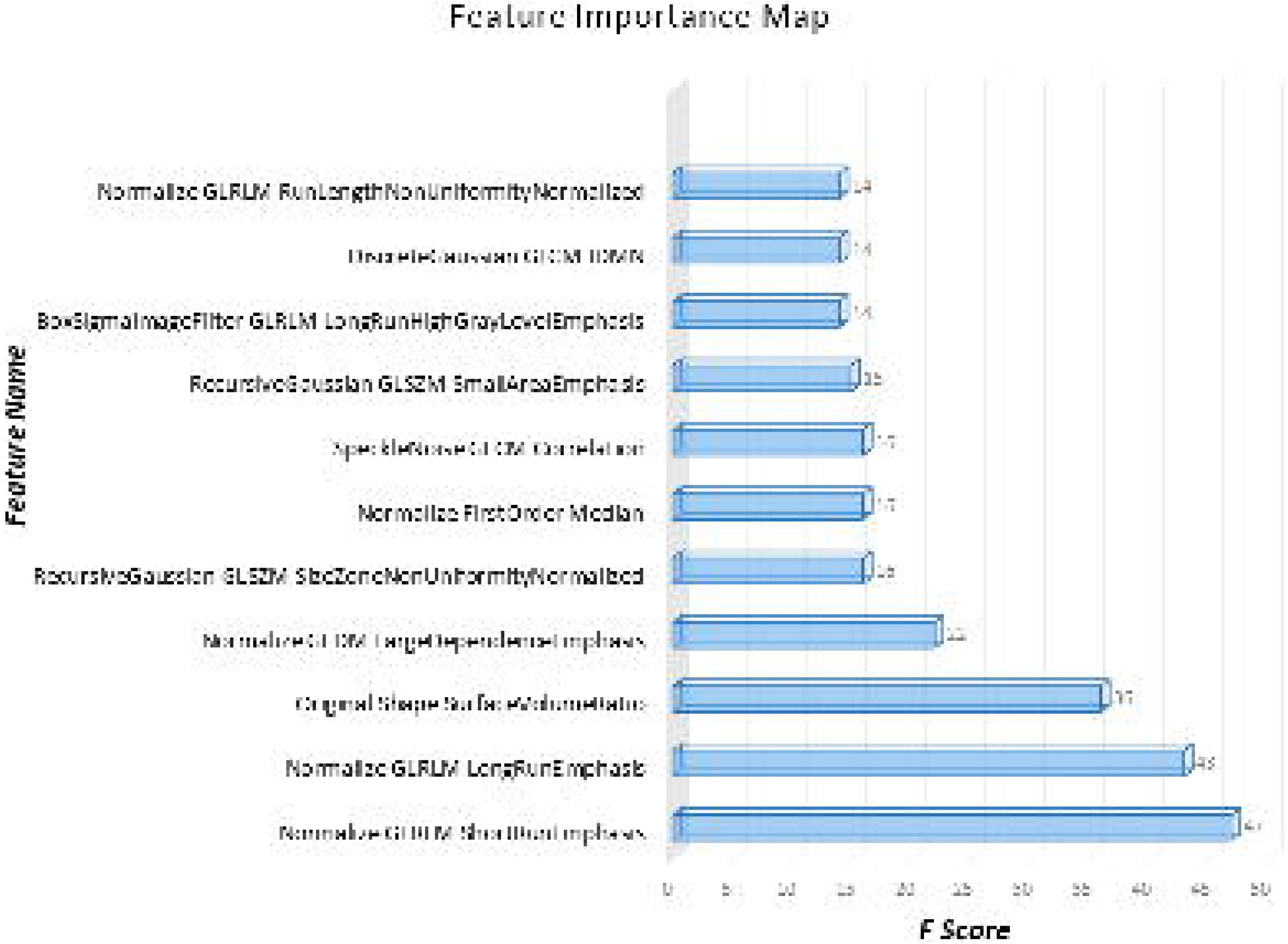
Radiomic features in order of their importance as plotted against the corresponding f-scores

Figure 4 demonstrates the class activation maps computed from CNNs for three patients. It was observed that in majority of the patients, the activations were more concentrated on the left SNc, which has been quantified and plotted in Figure 5. The activations on the left demonstrated a significant trend as they were greater than the right (p-value = 0.09). Similarly, for HCs, the activations on the left were more concentrated than on the right, however were not significant (p-value = 0.35).

**Figure 4:**
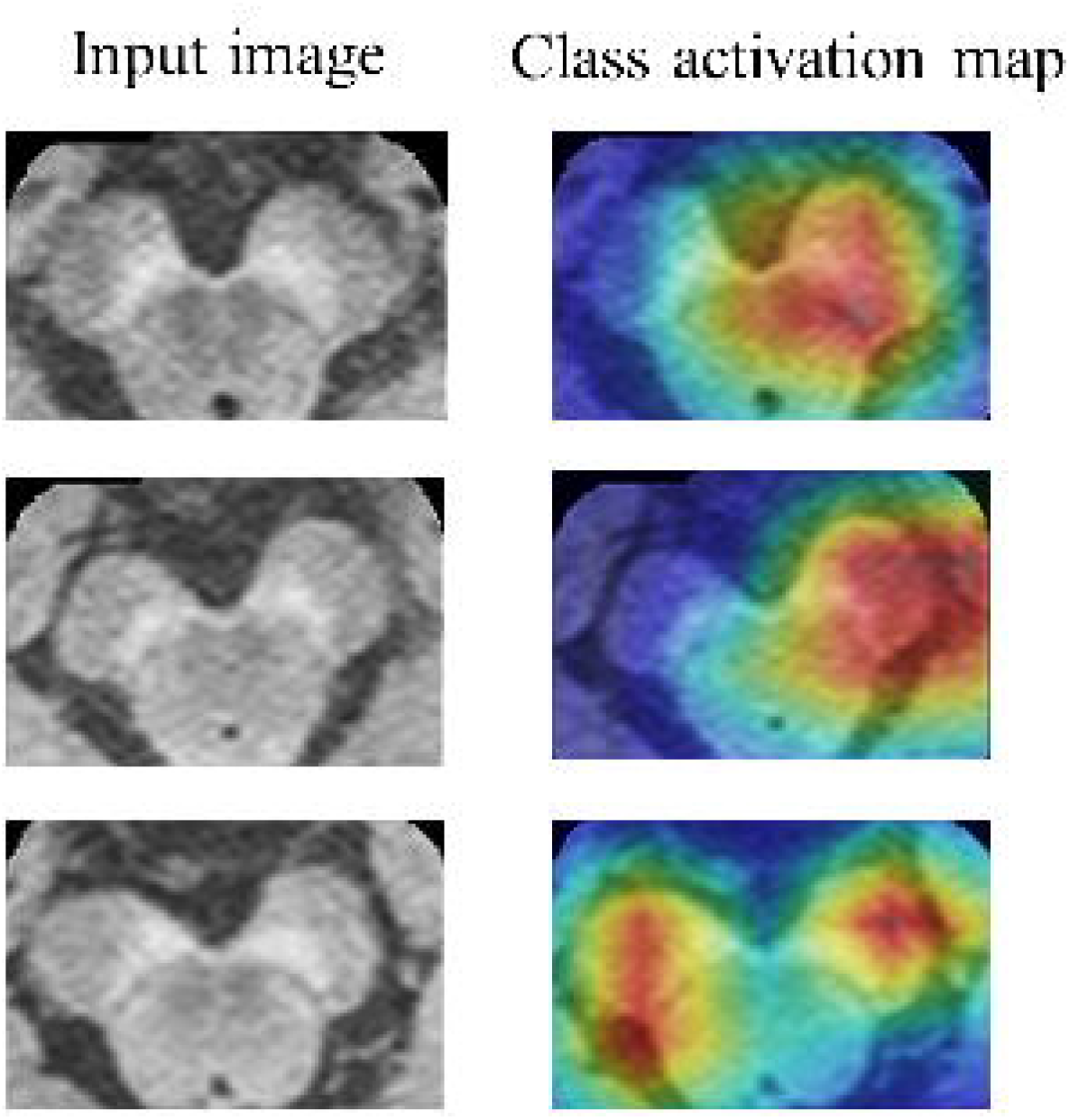
Examples of class Activation Maps of PD patients demonstrating that the SN area is highly activated while classifying PDs from Controls. In the first two subjects it can be observed that the left SNc is activated while in the third subject left and right SNc both are activated.

**Figure 5:**
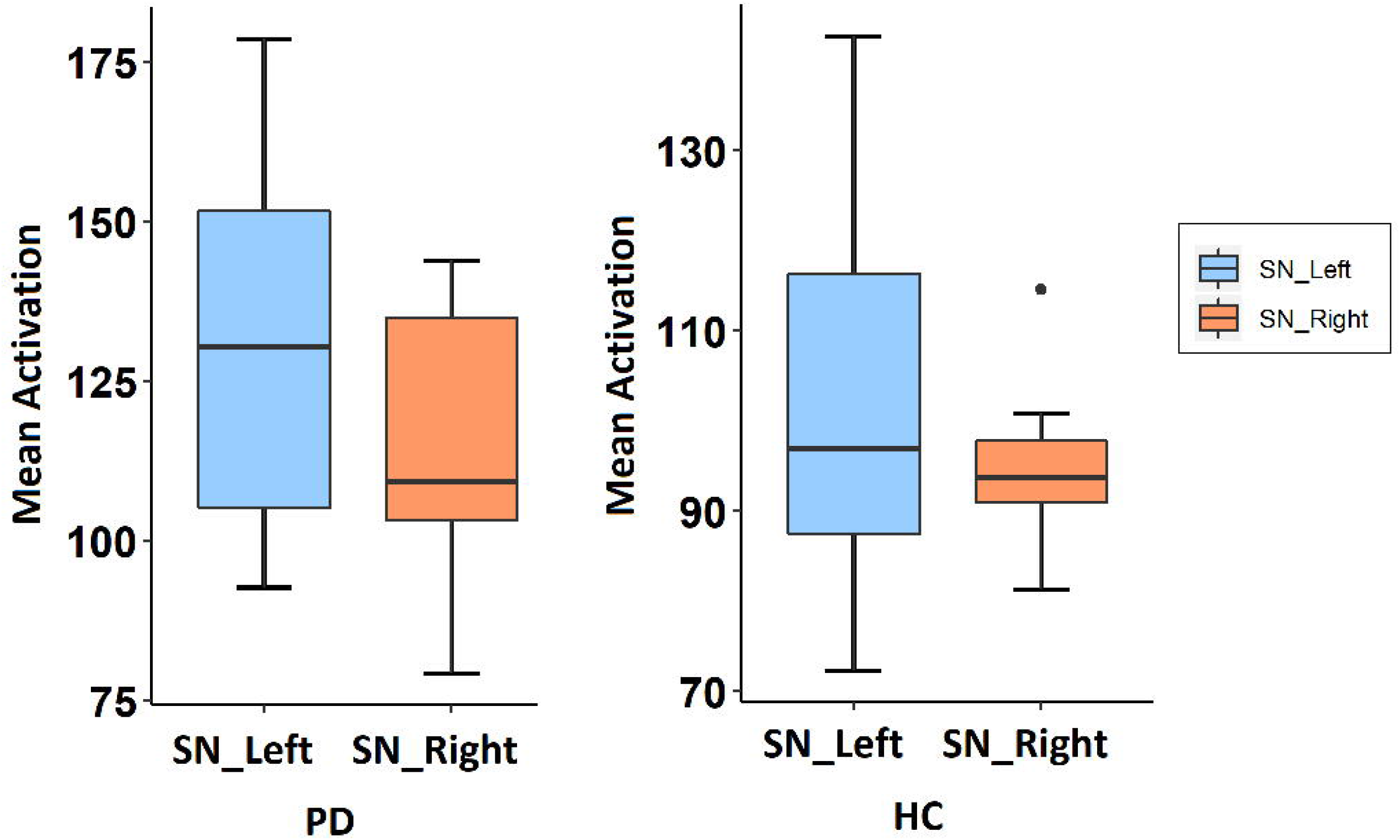
Boxplot demonstrating the asymmetry in activations computed from the CNNs.

## Discussion

This work presents a framework for creating predictive markers of PD using state-of-art CNNs based on NMS-MRI. We obtained a superior accuracy and demonstrated that our technique was better than contrast ratio-based predictions as well as predictions based off radiomic features computed from the SNc region, and therefore, may be utilized to support a PD diagnosis. Moreover, our CNN based classifier did not require extracting SNc borders as it employed a larger region of interest while the class activation maps illustrated the most important regions and with a significant trend in asymmetry which is in concurrence with the usual clinical picture of PD.

NMS-MRI has been frequently utilized and has demonstrated potential to differentiate PD from healthy controls and other parkinsonian and tremor disorders. Studies in NMS-MRI have shown that this sequence can be used to measure the concentration of the neuromelanin in SNc and lower signal intensities have been observed in patients with PD in comparison to healthy subjects (Ohtsuka et al., 2013; Prasad, Stezin, et al., 2018; Sasaki et al., 2006). However, these studies have been limited to contrast and volume features with group-wise uni-variate techniques used for analysis. Moreover, these features either require accurate boundaries of the SNc to be computed or placing manual ROIs to compute the contrast. We alleviated these setbacks in NMS-MRI analysis by employing an automated deep learning framework which once trained can be directly employed on every new patient facilitating a PD biomarker effortlessly, which may support in treatment planning and efficacy.

MR based machine learning for PD is not novel and multiple groups have made an effort in classifying PD (Table-1). However, none of these studies employ NMS-MRI, a powerful imaging technique for PD diagnosis as described earlier. Furthermore, most of these studies employ millions of features from single or multiple modalities which include voxel level features, on a petite sample size using SVMs that builds a hyperplane in the n-feature dimensional space. Even though such models achieve a good classification performance, it could be highly susceptible to over-fitting (Cherubini, Morelli, et al., 2014). To account for this, some studies extract the most effective features from voxel-based morphometry (VBM) analysis and use the mean values from significant clusters as features(Cherubini, Nistico, et al., 2014; Huppertz et al., 2016; Peran et al., 2018). Although this reduces the vulnerability to overfitting, extracting features off the population analysis may reduce the test accuracy as the pre-computed features from VBM may not be directly applicable to the test subjects. Unsupervised techniques such as self-organizing maps have also been employed on T1 MRIs(Singh & Samavedham, 2015). Our work is pioneering in terms of employing CNNs to classify PD, as CNNs can automatically extract relevant information from the images under consideration making the overall framework more applicable into a clinical setting. Such ability of CNNs in extracting discriminative features using local spatial coherence at various resolutions, is attractive, as it often accounts for better classification performance, especially on large datasets, when compared to a similar task on empirically drawn features such as CRs and/or Radiomic features in this case or VBM based clusters, diffusion measures etc. from earlier studies (Cherubini, Nistico, et al., 2014; Huppertz et al., 2016; Peran et al., 2018). We demonstrate this in Figure 2 where the AU-ROC of the CNN is superior to other techniques with manually extracted features. Such performance, however, comes with certain tradeoffs such as scarce information about the internal operation and behavior of these complex models which is especially important where the classes cannot be visually discerned (which perhaps includes majority classification problems in medical imaging). This is where our CAMs computed using the GAP layer play a crucial role in gaining insights into the functioning of the CNN as well as illustrating the most discriminative regions that can support clinical understanding. Figure 4 and 5 together demonstrate a trend in activations that are more concentrated in the left SNc than the right (p-value = 0.09) as has been shown before using manually extracted CRs (Prasad, Saini, et al., 2018). In controls, the left mean activation was higher than the right however with no trend for significance (p-value >0.3). Moreover, the mean activations in controls were lower than in patients suggesting that the model could effortlessly identify the regions which need to be highly weighted in PD subjects than in controls.

To compare our CNNs with other standard techniques we employed another novel method of extracting multiple radiomic features from the SNc region of interest. Radiomics, is a state-of-art technique more commonly employed in tumor classification problems. However, a recent study demonstrated the applicability of texture features in classifying PD using quantitative susceptibility maps and R2* maps (Li et al., 2019). In our case, RA-ML displayed superior cross-validation accuracy, however could not perform at the same level on the test data (Table 3). Figure 3 illustrates the most significant features that participated in the radiomics classification. These features involved grey level run length matrices that provide the size of homogenous runs for each grey level, non-uniformity measures, surface-volume ratios and grey level dependent matrix features that quantify the dependency of one voxel to another. These features overall captured the subtle changes in PD in the SNc that are revealed through NMS-MRI.

Although the NMS-MRI based CNN-DL classifier in the present study provides good accuracy in differentiating between PD and healthy controls, the clinical utility of this technique has to be determined by testing the ability of the classifier to differentiate between PD and other parkinsonian disorders such as progressive supranuclear palsy (PSP) and multiple system atrophy with predominant parkinsonism (MSA-P). Studies by Matsuura et al (Matsuura et al., 2013) and Ohtsuka et al (Ohtsuka et al., 2014) have demonstrated the utility of the NMS-MRI to differentiate between PD, MSA-P and PSP. Based on these observations we believe that our technique can be directly applied to differentiate various parkinsonian disorders.

It is important to note that the sample size of our study was relatively small; however, data augmentation techniques supported our classifier model(Ferreira et al.). Nevertheless, a larger sample size is certainly required to establish the utility of the method. Secondly, the regions of interest on the NMS-MRI that also included the brain stem were drawn manually, as input to the CNN, however, using deep learning-based techniques the boxed regions can be created automatically.

In summary, we introduced a novel computer aided PD diagnostic framework using the neuromelanin signal. Our findings show a superior accuracy using CNNs as compared to radiomics, while the underlying activation maps (Figure 4) confirm the involvement of SNc in the classification. This is commensurate with the idea of varying neuromelanin contrast on the SNc in the HCs and PDs thereby facilitating prediction of the underlying PD pathology.

## Acknowledgments

Symbiosis International University has received partial support from DST SERB (ECR/2016/000808) for setting up the computing facility.

