## Supplementary material for "Predictive markers for Parkinson’s disease using deep neural nets on neuromelanin sensitive MRI": Supplmentary data

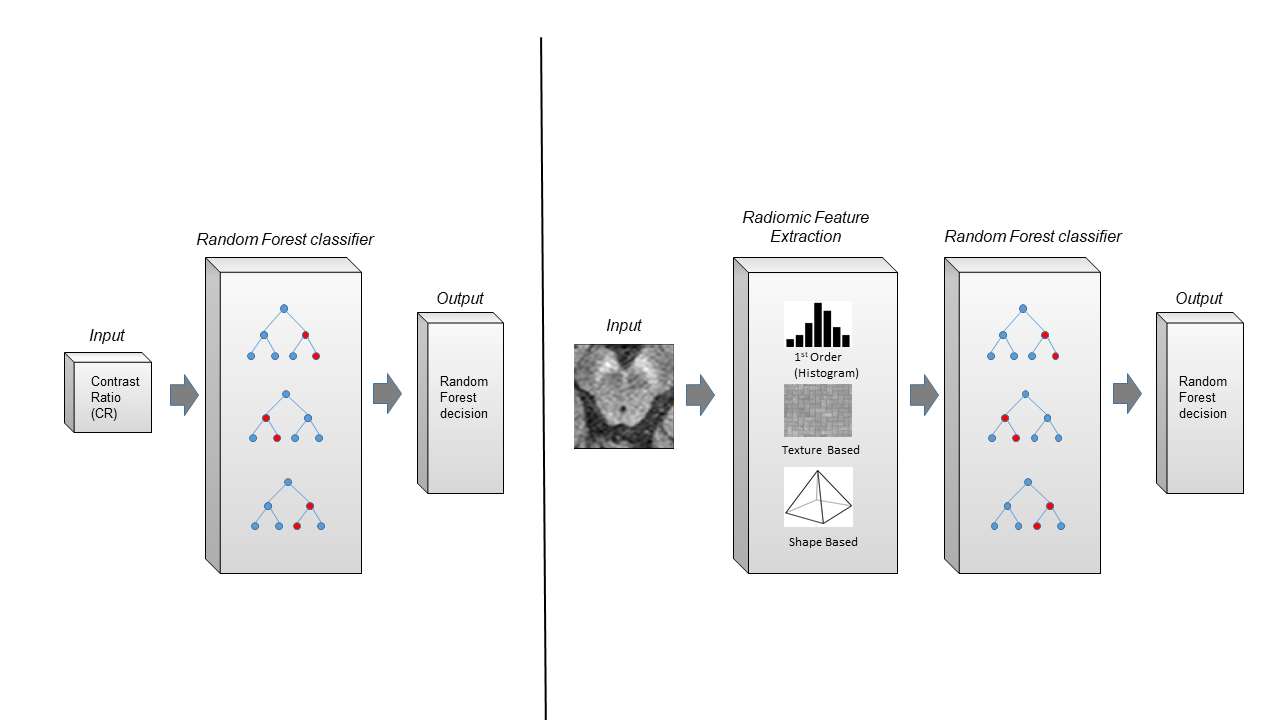

**Supplementary Figure 1: Figure displaying the procedure for CR-ML and RA-ML**

**Supplementary Table 1: Radiomics Features.**

First the feature extraction of the image without any filter takes place. Following this it undergoes 13 image filters which provide with 105 features per filter. Therefore, a total of 14*105 = 1470 features are computed per image. We describe the 105 features and the 13 filters employed in the below table.

#### Shape Based Features

This group of features includes descriptors of the three-dimensional size and shape of the ROI/ mask. These features are independent from the gray level intensity distribution in the ROI and are therefore only calculated on the non-derived image and mask.

A mesh is generated using marching cubes algorithm for the 3D MRI image.

Consider the following notations:

- *Nv* represents number of voxels in ROI
- *Nf* represents number of faces (triangles) defining the Mesh
- *V* represents the volume of the mesh in mm^3^
- *A* represents the surface area of the mesh in mm^2^

| S. No | Feature Name | Formula | Description |
| --- | --- | --- | --- |
| 1 | Volume | $V_{i}=\frac{O_{ai}\cdot\left( O_{bi}\times O_{ci} \right)}{6}$(1)  $V=\sum_{i=0}^{Nf} V_{i}$ (2) | The volume of the ROI is calculated from the triangle mesh of the ROI.  For each face i in the mesh, defined by points a_i_, b_i_ and c_i_, the (signed) volume V_f_ of the tetrahedron defined by that face and the origin of the image (O) is calculated in equation (1). We take the sum of all *V_i_* to compute *V.* |
| 2 | Sphericity | $sphericity=\frac{\sqrt[3]{36\pi V^{2}}}{A}$ | It is the measure of the roundness of the shape of the tumour relative to a sphere.  The value ranges from 0<*s**p**h**e**r**i**c**i**t**y*≤1, where a value of 1 indicates a perfect sphere. |
| 3 | Elongation | $elongation=\frac{\lambda_{minor}}{\lambda_{major}}$ | Elongation shows the relationship between the two largest principal components in ROI. *λ**_major_* and *λ**_minor_* are the lengths of the largest and second largest principal component axes. |
| 4 | Surface Volume Ratio | $Ratio=\frac{A}{V}$ | It is the ratio between the surface area and the volume.  A lower value indicates a more compact (sphere-like) shape. |
| 5 | Minor Axis | $minoraxis=4\sqrt{\lambda_{minor}}$ | This feature yields the second-largest axis length of the ROI-enclosing ellipsoid, where is the largest principal component axis. |
| 6 | Major Axis | $majoraxis=4\sqrt{\lambda_{major}}$ | This feature yield the largest axis length of the ROI-enclosing ellipsoid and is calculated using the largest principal component λ_major_. |
| 7 | Surface Area | $A_{i}=\frac{1}{2}\left\vert a_{i}b_{i}\times a_{i}c_{i} \right\vert$(1)   $A=\sum_{i=0}^{Nf} A_{i}$(2) | a*_i_*b*_i_* and a*_i_*c*_i_* are edges of the *i*^th^ triangle in the mesh, formed by vertices a*_i_*, b*_i_* and c*_i_*  First the surface area of each triangle is calculated in (1).  The total surface area is the sum of all calculated sub-areas. |
| 8 | Flatness | $flatness=\sqrt{\frac{\lambda_{least}}{\lambda_{major}}}$ | Flatness feature shows the relationship between the largest and smallest principal components in the ROI shape.  Here, *λ**_l_**_e_**_a_**_s_**_t_* and *λ**_m_**_a_**_j_**_o_**_r_* are the length of the largest and smallest principal component axes. |
| 9 | Least Axis | $leastaxis=4\sqrt{\lambda_{least}}$ | This feature yields the smallest axis length of the ROI-enclosing ellipsoid.  In case of a 2D segmentation, this value will be 0. |
| 10 | Maximum 2D Diameter Slice | Returns self.diameters[0] | The largest pair-wise Euclidean distance between tumour surface mesh vertices in the row-column (generally the axial) plane |
| 11 | Maximum 2D Diameter Column | Returns self.diameters[1] | It is defined as the largest pair-wise Euclidean distance between tumour surface mesh vertices in the row-slice (usually the coronal) plane. |
| 12 | Maximum 2D Diameter Row | Returns self.diameters[2] | It is defined as the largest pair-wise Euclidean distance between tumour surface mesh vertices in the column-slice (usually the sagittal) plane |
| 13 | Maximum 3D Diameter | Returns self.diameters[3] | The largest pair-wise Euclidean distance between the tumour surface mesh vertices. |

#### Gray Level Dependence Matrix (GLDM) Features

A Gray Level Dependence Matrix quantifies the gray level dependencies in an image. Gray level dependency is defined as the number of connected voxels within a distance *δ* that are dependent on the center voxel.

Consider the following notations:

- *Ng* represents the number of discreet intensity values in the image
- *Nd* represents the number of discreet dependency sizes in the image
- *P(i, j)* represents the dependence matrix
- *p(i, j)* represents the normalized dependence matrix, defined as$p\left( i,j \right)=\frac{P\left( i,j \right)}{Nz}$
- *Nz* represents the number of dependency zones in the image, which is equal to

| S. No | Feature Name | Formula | Description |
| --- | --- | --- | --- |
| 14 | Gray Level Variance | $GLV=\sum_{i=0}^{Ng} \sum_{j=0}^{Nd} p\left( i,j \right)\left( i-\mu\right)^{2}$  where:  $\mu=\sum_{i=1}^{Ng} \sum_{j=1}^{Nd} ip\left( i,j \right)$ | Measures the variance in grey level in the image. |
| 15 | Low Gray Level Emphasis | $LGLE=\frac{\sum_{i=0}^{Ng} \sum_{j=0}^{Nd} \left( \frac{p\left( i,j \right)}{i^{2}} \right)}{Nz}$ | Measures the distribution of low gray level values.  A higher value indicates a greater concentraton of low gray level values in the image. |
| 16 | High Gray Level Emphasis | $\frac{\sum_{i=1}^{Ng} \sum_{j=1}^{Nd} P\left( i,j \right)i^{2}}{Nz}$ | Measures the joint distribution of small dependence with lower gray-level values. |
| 17 | Dependence Entropy | $-\sum_{i=0}^{Ng} \sum_{j=0}^{Nd} p\left( i,j \right)log\left( p\left( i,j \right)+\epsilon\right)$ | Measures the randomness /variability of dependence throughout the image. |
| 18 | Dependence Non Uniformity | $DN=\frac{\sum_{j=1}^{Nd} \left( \sum_{i=1}^{Ng} P\left( i,j \right) \right)^{2}}{Nz}$ | Measures the similarity of dependence throughout the image. |
| 19 | Dependence Non Uniformity Normalized | $DNN=\frac{\sum_{j=1}^{Nd} \left( \sum_{i=1}^{Ng} P\left( i,j \right) \right)^{2}}{{Nz}^{2}}$ | Similar to Dependence Non Uniformity, however contains the normalized result. |
| 20 | Gray Level Non Uniformity | $GLN=\frac{\sum_{i=0}^{Ng} \left( \sum_{j=1}^{Nd} P\left( i,j \right) \right)^{2}}{Nz}$ | Measures the similarity of gray level intensity values in the image. |
| 21 | Small Dependence Emphasis | $SDE=\frac{\sum_{i=1}^{Ng} \sum_{j=1}^{Nd} \left( \frac{P\left( i,j \right)}{i^{2}} \right)}{Nz}$ | It is a measure of the distribution of small dependencies.   A high value is indicative of small dependence and less homogeneous textures. |
| 22 | Large Dependence Emphasis | $LDE=\frac{\sum_{i=1}^{Ng} \sum_{j=1}^{Nd} P\left( i,j \right)j^{2}}{Nz}$ | It is a measure of the distribution of large dependencies.  A high value is indicative of large dependence and more homogeneous textures. |
| 23 | Small Dependence Low Gray Level Emphasis | $\frac{\sum_{i=1}^{Ng} \sum_{j=1}^{Nd} \frac{P\left( i,j \right)}{i^{2}j^{2}}}{N_{z}}$ | It measures the joint distribution of small dependence with lower gray-level values. |
| 24 | Small Dependence High Gray Level Emphasis | $\frac{\sum_{i=1}^{Nd} \sum_{j=1}^{Nd} \frac{P\left( i,j \right)i^{2}}{j^{2}}}{Nz}$ | It measures the joint distribution of small dependence with higher gray-level values. |
| 25 | Large Dependence Low Gray Level Emphasis | $\frac{\sum_{i=1}^{Ng} \sum_{j=1}^{Nd} \frac{P\left( i,j \right)j^{2}}{i^{2}}}{N_{z}}$ | It measures the joint distribution of large dependence with lower gray-level values. |
| 26 | Large Dependence High Gray Level Emphasis | $\frac{\sum_{i=1}^{Ng} \sum_{j=1}^{Nd} P\left( i,j \right)i^{2}j^{2}}{Nz}$ | It measures the joint distribution of large dependence with higher gray-level values. |
| 27 | Dependence Variance | $DV=\sum_{i=1}^{Ng} \sum_{j=1}^{Nd} p\left( i,j \right)\left( j-\mu\right)^{2}$ $\mathrm{where}\mu=\sum_{i=1}^{Ng} \sum_{j=1}^{Nd} jp\left( i,j \right)$ | Measures the variance in dependence size in the image. |

#### Gray Level Co-occurrence Matrix (GLCM) Features

A Gray Level Co-occurrence Matrix (GLCM) of size N_g_×N_g_N_g_×N_g_ describes the second-order joint probability function of an image region constrained by the mask and is defined as P(i,j|δ,θ)P(i,j|δ,θ). The (i,j)th(i,j)th element of this matrix represents the number of times the combination of levels iiand jj occur in two pixels in the image, that are separated by a distance of δδ pixels along angle θθ. The distance δδ from the center voxel is defined as the distance according to the infinity norm. For δ=1δ=1, this results in 2 neighbors for each of 13 angles in 3D (26-connectivity) and for δ=2δ=2 a 98-connectivity (49 unique angles).

Consider the following:

- ϵ represents an arbitrarily small positive number (≈2.2×10−16≈2.2×10−16)
- *P(i,j)* represents the co-occurence matrix for an arbitrary δδ and θθ
- *p(i,j)* represents the normalized co-occurence matrix and equal to P(i,j)∑P(i,j)P(i,j)∑P(i,j)
- *N_g_* represents the number of discrete intensity levels in the image
- $p_{x}\left( i \right)=\sum_{j=1}^{Ng} P\left( i,j \right)$represents the marginal row probabilities
- $p_{y}\left( j \right)=\sum_{j=1}^{Ng} P\left( i,j \right)$ represents the marginal column probabilities
- $\mu_{x}$ represents the mean gray level intensity of $p_{x}$ and defined as $\mu_{x}=\sum_{i=1}^{Ng} p_{x}\left( i \right)i$
- $\mu_{y}$ represents the mean gray level intensity of $p_{y}$ and defined as $\mu_{y}=\sum_{j=1}^{Ng} p_{y}\left( j \right)j$
- $\sigma_{x}$represents the standard deviation of $p_{x}$
- $\sigma_{y}$ represent the standard deviation of $p_{y}$
- $p_{x+y}\left( k \right)=\sum_{i=1}^{Ng} \sum_{j=1}^{Ng} p\left( i,j \right)$, where $k=i+j$ and $k=2,3,\ldots.,2Ng$
- $p_{x-y}\left( k \right)=\sum_{i=1}^{Ng} \sum_{j=1}^{Ng} p\left( i,j \right)$, where $\left| i-j \right|=k$, and $k=0,1,\ldots,Ng-1$
- $HX=-\sum_{i=1}^{Ng} p_{x}\left( i \right){log}_{2}\left( p_{x}\left( i \right)+\epsilon\right)$represents the entropy of $p_{x}$
- $HY=-\sum_{j=1}^{Ng} p_{y}\left( j \right){log}_{2}\left( p_{y}\left( j \right)+\epsilon\right)$represents the entropy of $p_{y}$
- $HXY=-\sum_{i=1}^{Ng} \sum_{j=1}^{Ng} p\left( i,j \right){log}_{2}\left( p\left( i,j \right)+\epsilon\right)$ be the entropy of $p\left( i,j \right)$
- $HXY1=-\sum_{i=1}^{Ng} \sum_{j=1}^{Ng} p\left( i,j \right){log}_{2}\left( p_{x}\left( i \right)p_{y}\left( j \right)+\epsilon\right)$
- $HXY2=-\sum_{i=1}^{Ng} \sum_{j=1}^{Ng} p_{x}\left( i \right)p_{y}\left( j \right){log}_{2}\left( p_{x}\left( i \right)p_{y}\left( j \right)+\epsilon\right)$

| Sr. No | Feature Name | Formula | Description |
| --- | --- | --- | --- |
| 28 | Joint Average | $\mu x=\sum_{i=1}^{Ng} \sum_{j=1}^{Ng} p\left( i,j \right)i$ | Joint average- *μ**x* returns the mean gray level intensity of the *i* distribution. |
| 29 | Autocorrelation | $\sum_{i=1}^{Ng} \sum_{j=1}^{Ng} p\left( i,j \right)ij$ | Autocorrelation is a measure of the magnitude of the fineness and coarseness of texture. |
| 30 | Cluster Prominence | $\sum_{i=1}^{Ng} \sum_{j=1}^{Ng} \left( i+j+\mu x+\mu\right)^{4}p\left( i,j \right)$ | Cluster Prominence is a measure of the skewness and asymmetry of the GLCM.   A higher value implies more asymmetry about the mean. A lower value implies less variation about the mean. |
| 31 | Cluster Shade | $\sum_{i=1}^{Ng} \sum_{j=1}^{Ng} \left( i+j-\mu x-\mu\right)^{3}p\left( i,j \right)$ | Cluster Shade is a measure of the skewness and uniformity of the GLCM.  A higher cluster shade implies greater asymmetry about the mean. |
| 32 | Cluster Tendency | $\sum_{i=1}^{Ng} \sum_{j=1}^{Ng} \left( i+j-\mu x-\mu\right)^{2}p\left( i,j \right)$ | Cluster Tendency is a measure of groupings of voxels with similar gray-level values. |
| 33 | Sum Average | $\sum_{k=2}^{2Ng} p_{x+y}\left( k \right)k$ | Sum Average measures the relationship between occurrences of pairs with lower intensity values and occurrences of pairs with higher intensity values. |
| 34 | Difference Average | $\sum_{k=0}^{Ng-1} kp_{x-y}\left( k \right)$ | Difference Average measures the relationship between occurrences of pairs with similar intensity values and occurrences of pairs with differing intensity values. |
| 35 | Joint Energy | $\sum_{i=1}^{Ng} \sum_{j=1}^{Ng} \left( p\left( i,j \right) \right)^{2}$ | Energy is a measure of homogeneous patterns in the image.  A greater Energy implies that there are more instances of intensity value pairs in the image that neighbor each other at higher frequencies. |
| 36 | Joint Entropy | $-\sum_{i=1}^{Ng} \sum_{j=1}^{Ng} p\left( i,j \right){log}_{2}\left( p\left( i,j \right)+\epsilon\right)$ | Joint entropy is a measure of the randomness/ variability in neighbourhood intensity values. |
| 37 | Difference Entropy | $\sum_{k=0}^{Ng-1} p_{x-y}\left( k \right){log}_{2}\left( p_{x-y}\left( k \right)+\epsilon\right)$ | Difference Entropy is a measure of the randomness/ variability in neighbourhood intensity value differences. |
| 38 | Difference Variance | $\sum_{k=0}^{Ng-1} \left( k-DA \right)^{2}p_{x-y}\left( k \right)$ | Difference Variance is a measure of heterogeneity that places higher weights on differing intensity level pairs that deviate more from the mean. |
| 39 | Inverse Variance | $\sum_{k=1}^{Ng-1} \frac{p_{x-y}\left( k \right)}{k^{2}}$ | Inverse Variance is a method in which each weight is measured by the inverse of the variance of the effect size. |
| 40 | Maximum Probability | $max\left( p\left( i,j \right) \right)$ | Maximum Probability is occurrences of the most predominant pair of neighboring intensity values. |
| 41 | Contrast | $\sum_{i=1}^{Ng} \sum_{j=1}^{Ng} \left( i-j \right)^{2}p\left( i,j \right)$ | Contrast is a measure of the local intensity variation, favouring values away from the diagonal (i=j)(i=j).  A larger value correlates with a greater disparity in intensity values among neighbouring voxels. |
| 42 | Correlation | $\frac{\sum_{i=1}^{Ng} \sum_{j=1}^{Ng} p\left( i,j \right)ij-\mu_{x}-\mu_{y}}{\sigma_{x}\left( i \right)\sigma_{y}\left( j \right)}$ | Correlation is a value between 0 (uncorrelated) and 1 (perfectly correlated) showing the linear dependency of gray level values to their respective voxels in the GLCM. |
| 43 | Sum of Squares | $\sum_{i=1}^{Ng} \sum_{j=1}^{Ng} \left( i-\mu_{x} \right)^{2}p\left( i,j \right)$ | Sum of Squares or Variance is a measure in the distribution of neighboring intensity level pairs about the mean intensity level in the GLCM. |
| 44 | ID – Inverse Difference | $\sum_{k=0}^{Ng-1} \frac{p_{x-y}\left( k \right)}{1+k}$ | ID is another measure of the local homogeneity of an image. It is also known as Homogeneity 1.   With more uniform gray levels, the denominator will remain low, resulting in a higher overall value. |
| 45 | IDN – Inverse Difference Normalized | $\sum_{k=0}^{Ng-1} \frac{p_{x-y}\left( k \right)}{1+\frac{k}{Ng}}$ | Inverse Difference Normalized is another measure of the local homogeneity of an image.  IDN normalizes the difference between the neighboring intensity values by dividing over the total number of discrete intensity values. |
| 46 | IDM – Inverse Difference Moment | $\sum_{k=0}^{Ng-1} \frac{p_{x-y}\left( k \right)}{1+k^{2}}$ | IDM is a measure of the local homogeneity of an image. It is also known as Homogeneity 2.  IDM weights are the inverse of the Contrast weights (decreasing exponentially from the diagonal *i=j* in the GLCM). |
| 47 | IDMN – Inverse Difference Moment Normalized | $\sum_{k=0}^{Ng-1} \frac{p_{x-y}\left( k \right)}{1+\frac{k^{2}}{N_{g}^{2}}}$ | IDMN is a measure of the local homogeneity of an image.   IDMN normalizes the square of the difference between the neighbouring intensity values by dividing over the square of the total number of discrete intensity values. |
| 48 | Sum Entropy | $\sum_{k=2}^{2Ng} p_{x+y}\left( k \right){log}_{2}\left( p_{x+y}\left( k \right)+\epsilon\right)$ | Sum Entropy is a sum of neighborhood intensity value differences. |
| 49 | IMC1 – Informational Measure of Correlation 1 | $\frac{-I\left( x,y \right)}{max\{HX,HY\}}$    $\frac{HXY-HXY1}{max\{HX,HY\}}$ | IMC1 assesses the correlation between the probability distributions of *i* and *j* (quantifying the complexity of the texture), using mutual information *I(x, y)*  In the case where the distributions are independent, there is no mutual information and the result will therefore be 0. In the case of uniform distribution with complete dependence, mutual information will be equal to *log2 (Ng)*. |
| 50 | IMC2 – Informational Measure of Correlation 2 | $\sqrt{1-e^{-2\left( HXY2-HXY \right)}}$ | IMC2 also assesses the correlation between the probability distributions of *i* and *j* (quantifying the complexity of the texture). |

#### First Order Statistical Features

First-order statistics describe the distribution of voxel intensities within the image region defined by the mask through commonly used and basic metrics.

Consider the following:

- Let **X** be a set of *Np* voxels included in the ROI.
- Let **P*(i)*** be the first order histogram with ***Ng*** discrete intensity levels, where ***Ng*** is the number of non-zero bins, equally spaced from 0 with a width defined in the **binWidth** parameter.
- Let ***p(i)*** be the normalized first order histogram and equal to $\frac{P\left( i \right)}{Np}$

| S. No. | Feature Name | Formula | Description |
| --- | --- | --- | --- |
| 51 | Energy | $\sum_{i=1}^{Np} \left( X\left( i \right)+c \right)^{2}$ | Energy is a measure of the magnitude of voxel values in an image.   A larger values implies a greater sum of the squares of these values.  Here, *c* is optional value, defined by *voxelArrayShift*, which shifts the intensities to prevent negative values in X. This ensures that voxels with the lowest gray values contribute the least to Energy, instead of voxels with gray level intensity closest to 0. |
| 52 | Total Energy | $V_{voxel}\sum_{i=1}^{Np} \left( X\left( i \right)+c \right)^{2}$ | Total Energy is the value of Energy feature scaled by the volume of the voxel $\left( V \vert voxel \right)$ in cubic mm.  Here, cc is optional value, defined by *voxelArrayShift*, which shifts the intensities to prevent negative values in X |
| 53 | Entropy | $-\sum_{i=1}^{Np} p\left( i \right){log}_{2}\left( p\left( i \right)+\epsilon\right)$ | Entropy specifies the uncertainty/ randomness in the image values. It measures the average amount of information required to encode the image values.  $\epsilon$ is an arbitrarily small positive number (≈2.2×10−16≈2.2×10−16). |
| 54 | Maximum | $maximum=max\left( X \right)$ | The maximum gray level intensity within the ROI. |
| 55 | Minimum | $minimum=min\left( X \right)$ | The minimum gray level intensity within the ROI. |
| 56 | 10^th^ Percentile | *numpy.percentile(VoxelArray, 10):* Calls function for the 10^th^ percentile of the voxels. | Returns the 10^th^ percentile of X. |
| 57 | 90^th^ Percentile | *numpy.percentile(VoxelArray, 90):* Calls function for the 90^th^ percentile of the voxels. | Returns the 90^th^ percentile of X. |
| 58 | Mean | $\frac{1}{Np}\sum_{i=1}^{Np} X\left( i \right)$ | Mean gives the average gray level intensity within the ROI. |
| 59 | Median | *numpy.median(VoxelArray)* Calls the function to calculate median of the voxels. | The median gray level intensity within the ROI. |
| 60 | Inter-quartile Range | $IQRange=P_{75}-P_{25}$ | Here $P_{75}$ and $P_{25}$ are the 75^th^ and 25^th^ percentile of the image array respectively. |
| 61 | Range | $range=max\left( X \right)-min\left( X \right)$ | This refers to the range of gray values in the ROI. |
| 62 | Mean Absolute Deviation | $\frac{1}{Np}\sum_{i=1}^{Np} \left\vert X\left( i \right)-\acute{X} \right\vert$ | Mean Absolute Deviation is the mean distance of all intensity values from the Mean Value of the image array. |
| 63 | Robust Mean Absolute Deviation | $\frac{1}{N_{10-90}}\sum_{i=1}^{N_{10-90}} \left\vert X_{10-90}\left( i \right)-\acute{X}_{10-90} \right\vert$ | Robust Mean Absolute Deviation is the mean distance of all intensity values from the Mean Value calculated on the subset of image array with gray levels in between, or equal to the 10^th^ and 90^th^ percentile. |
| 64 | Root Mean Squared | $\sqrt{\frac{1}{Np}\sum_{i=1}^{Np} \left( X\left( i \right)+c \right)^{2}}$ | RMS is the square-root of the mean of all the squared intensity values. It is another measure of the magnitude of the image values.  Here, *c* is optional value, defined by *voxelArrayShift*, which shifts the intensities to prevent negative values in X.  This feature is volume-confounded; a larger value of *c* increases the effect of volume-confounding. |
| 65 | Skewness | $skewness=\frac{\mu_{3}}{\sigma^{3}}$ $\frac{\frac{1}{Np}\sum_{i=1}^{Np} \left( X\left( i \right)-\acute{X} \right)^{3}}{\left( \sqrt{\frac{1}{Np}\sum_{i=1}^{Np} \left( X\left( i \right)-\acute{X} \right)^{2}} \right)^{3}}$ | Skewness measures the asymmetry of the distribution of values about the Mean value.   Depending on where the tail is elongated and the mass of the distribution is concentrated, this value can be positive or negative.  Here $\mu_{3}$ is the 3^rd^ central moment. |
| 66 | Kurtosis | $kurtosis=\frac{\mu_{4}}{\sigma^{4}}$ $\frac{\frac{1}{Np}\sum_{i=1}^{Np} \left( X\left( i \right)-\acute{X} \right)^{4}}{\left( \frac{1}{Np}\sum_{i=1}^{Np} \left( X\left( i \right)-\acute{X} \right)^{2} \right)^{2}}$ | Kurtosis is a measure of the ‘peakedness’ of the distribution of values in the image ROI.  A higher kurtosis implies that the mass of the distribution is concentrated towards the tail(s) rather than towards the mean. A lower kurtosis implies the reverse: that the mass of the distribution is concentrated towards a spike near the Mean value.  Here $\mu_{4}$ is the 4^th^ central moment. |
| 67 | Variance | $variance=\sigma^{2}$ $\frac{1}{Np}\sum_{i=1}^{Np} \left( X\left( i \right)-\acute{X} \right)^{2}$ | Variance is the mean of the squared distances of each intensity value from the Mean value.   It is a measure of the spread of the distribution about the mean. |
| 68 | Uniformity | $uniformity=\sum_{i=1}^{Ng} {p\left( i \right)}^{2}$ | Uniformity is a measure of the sum of the squares of each intensity value.  It is a measure of the homogeneity of the image array, where a greater uniformity implies a greater homogeneity or a smaller range of discrete intensity values. |

#### Gray Level Run Length Matrix (GLRLM) Features

A Gray Level Run Length Matrix (GLRLM) quantifies gray level runs, which are defined as the length in number of pixels, of consecutive pixels that have the same gray level value. In a gray level run length matrix ***P****(i,j|θ)*, the *(i,j)*^th^ element describes the number of runs with gray level *i* and length *j* occur in the image (ROI) along angle *θ*.

Let:

- *Ng* be the number of discreet intensity values in the image.
- *Nr* be the number of discreet run lengths in the image.
- *Np* be the number of voxels in the image
- *Nz(θ)* be the number of runs in the image along angle θ, which is equal to $\sum_{i=1}^{Ng} \sum_{j=1}^{Nr} P\left( i,j|\theta\right)$ and $1\leq N_{z}\left( \theta\right)$
- ***P****(i,j|θ)* be the run length matrix for an arbitrary direction *θ*
- *p(i,j|θ)* be the normalized run length matrix, defined as $p\left( i,j | \theta\right)=\frac{P\left( i,j|\theta\right)}{N_{z}\left( \theta\right)}$

| S. No. | Feature Name | Formula | Description |
| --- | --- | --- | --- |
| 69 | Short Run Emphasis | $\frac{\sum_{i=1}^{Ng} \sum_{j=1}^{Nr} \frac{P\left( i,j\vert\theta\right)}{j^{2}}}{N_{z}\left( \theta\right)}$ | It is a measure of the distribution of short run lengths, with a greater value indicative of shorter run lengths and more fine textural textures. |
| 70 | Long Run Emphasis | $\frac{\sum_{i=1}^{Ng} \sum_{j=1}^{Nr} P\left( i,j\vert\theta\right)j^{2}}{N_{z}\left( \theta\right)}$ | It is a measure of the distribution of long run lengths, with a greater value indicative of longer run lengths and more coarse structural textures. |
| 71 | Gray Level Non Uniformity | $\frac{\sum_{i=1}^{Ng} \left( \sum_{j=1}^{Nr} P\left( i,j\vert\theta\right) \right)^{2}}{N_{z}\left( \theta\right)}$ | GLN measures the similarity of gray-level intensity values in the image, where a lower GLN value correlates with a greater similarity in intensity values. |
| 72 | Gray Level Non Uniformity Normalized (GLNN) | $\frac{\sum_{i=1}^{Ng} \left( \sum_{i=1}^{Ng} P\left( i,j\vert\theta\right) \right)^{2}}{N_{z}\left( \theta\right)}$ | It measures the similarity of gray-level intensity values in the image, where a lower GLNN value correlates with a greater similarity in intensity values.  This is the normalized version of the GLN formula. |
| 73 | Run Length Non Uniformity (RLN) | $\frac{\sum_{j=1}^{Nr} \left( \sum_{i=1}^{Ng} P\left( i,j\vert\theta\right) \right)^{2}}{N_{z}\left( \theta\right)}$ | It measures the similarity of run lengths throughout the image, with a lower value indicating more homogeneity among run lengths in the image. |
| 74 | Run Length Non Uniformity N`ormalized (RLNN) | $\frac{\sum_{j=1}^{Nr} \left( \sum_{i=1}^{Ng} P\left( i,j\vert\theta\right) \right)^{2}}{{N_{z}\left( \theta\right)}^{2}}$ | It measures the similarity of run lengths throughout the image, with a lower value indicating more homogeneity among run lengths in the image. This is the normalized version of the RLN formula. |
| 75 | Run Percentage | $\frac{N_{z}\left( \theta\right)}{N_{p}}$ | RP measures the coarseness of the texture by taking the ratio of number of runs and number of voxels in the ROI.  Values are in range$\frac{1}{N_{p}}<RP<1$; with higher values indicating a larger portion of the ROI consists of short runs (indicates a more fine texture). |
| 76 | Gray Level Variance | $\sum_{i=1}^{Ng} \sum_{j=1}^{Nr} p\left( i,j \vert\theta\right)\left( i-\mu\right)^{2}$ $where,$ $\mu=\sum_{i=1}^{Ng} \sum_{j=1}^{Nr} p\left( i,j \vert\theta\right)i$ | GLV measures the variance in gray level intensity for the runs. |
| 77 | Run Variance | $\sum_{i=1}^{Ng} \sum_{j=1}^{Nr} p\left( i,j \vert\theta\right)\left( j-\mu\right)^{2}$ | RV is a measure of the variance in runs for the run lengths. |
| 78 | Run Entropy | $\sum_{i=1}^{Ng} \sum_{j=1}^{Nr} p\left( i,j \vert\theta\right){log}_{2}$ | It measures the uncertainty/ randomness in the distribution of run lengths and gray levels.   A higher value indicates more heterogeneity in the texture patterns.  Here ϵ is an arbitrarily small positive number $\left( \approx2.2\times{10}^{-16} \right)$ |
| 79 | Low Gray Level Run Emphasis | $\frac{\sum_{i=1}^{Ng} \sum_{j=1}^{Nr} \frac{P\left( i,j\vert\theta\right)}{i^{2}}}{N_{z}\left( \theta\right)}$ | It measures the distribution of low gray-level values, with a higher value indicating a greater concentration of low gray-level values in the image. |
| 80 | High Gray Level Run Emphasis | $\frac{\sum_{i=1}^{Ng} \sum_{j=1}^{Nr} P\left( i,j\vert\theta\right)i^{2}}{N_{z}\left( \theta\right)}$ | It measures the distribution of the higher gray-level values.  A higher value indicates a greater concentration of high gray-level values in the image. |
| 81 | Short Run Low Gray Level Emphasis | $\frac{\sum_{i=1}^{Ng} \sum_{j=1}^{Nr} \frac{P\left( i,j\vert\theta\right)}{i^{2}j^{2}}}{N_{z}\left( \theta\right)}$ | It measures the joint distribution of shorter run lengths with lower gray-level values. |
| 82 | Short Run High Gray Level Emphasis | $\frac{\sum_{i=1}^{Ng} \sum_{j=1}^{Nr} \frac{P\left( i,j\vert\theta\right)i^{2}}{j^{2}}}{N_{z}\left( \theta\right)}$ | It measures the joint distribution of shorter run lengths with higher gray-level values. |
| 83 | Long Run Low Gray Level Emphasis | $\frac{\sum_{i=1}^{Ng} \sum_{j=1}^{Nr} \frac{P\left( i,j\vert\theta\right)j^{2}}{i^{2}}}{N_{z}\left( \theta\right)}$ | It measures the joint distribution of long run lengths with lower gray-level values. |
| 84 | Long Run High Gray Level Emphasis | $\frac{\sum_{i=1}^{Ng} \sum_{j=1}^{Nr} P\left( i,j\vert\theta\right)i^{2}j^{2}}{N_{z}\left( \theta\right)}$ | It measures the joint distribution of long run lengths with higher gray-level values. |

#### Gray Level Size Zone Matrix (GLSZM) Features

A Gray Level Size Zone (GLSZM) quantifies the gray level zones in an image. A gray level zone is defined as the number of connected voxels that share the same gray level intensity. A voxel is considered connected if the distance is 1 according to the infinity norm (26-connected region in a 3D, 8-connected region in 2D).

In a gray level size zone matrix *P(i,j)*, the *(i,j)*^th^ element equals to the number of zones with gray level *i* and size *j* appear in image. Contrary to GLCM and GLRLM, the GLSZM is rotation independent, with only one matrix calculated for all directions in the ROI.

Let:

- *Ng* be the number of discreet intensity values in the image
- *Ns* be the number of discreet zone sizes in the image
- *Np* be the number of voxels in the image
- *Nz* be the number of zones in the ROI, which is equal to
  $\sum_{i=1}^{Ng} \sum_{j=1}^{Ns} P\left( i,j \right)$ and $1\leq N_{z}\leq N_{p}$
- *P(i,j)* be the size zone matrix
- *p(i,j)* be the normalized size zone matrix, defined as $p\left( i,j \right)=\frac{P\left( i,j \right)}{N_{z}}$

| Sr. No | Feature Name | Formula | Description |
| --- | --- | --- | --- |
| 85 | Small Area Emphasis | $\frac{\sum_{i=1}^{Ng} \sum_{j=1}^{Ns} \frac{P\left( i,j \right)}{j^{2}}}{Nz}$ | It is a measure of the distribution of small size zones.  A high value indicates smaller size zones and more fine textures. |
| 86 | Large Area Emphasis | $\frac{\sum_{i=1}^{Ng} \sum_{j=1}^{Ns} P\left( i,j \right)j^{2}}{Nz}$ | It is a measure of the distribution of large area size zones.  A high value indicates large size zones and more coarse textures. |
| 87 | Gray Level Non Uniformity (GLN) | $\frac{\sum_{i=1}^{Ng} \left( \sum_{j=1}^{Ns} P\left( i,j \right) \right)^{2}}{Nz}$ | It measures the variability of gray-level intensity values in the image.  A low value indicates more homogeneity in intensity values. |
| 88 | Gray Level Non Uniformity Normalized (GLNN) | $\frac{\sum_{i=1}^{Ng} \left( \sum_{j=1}^{Ns} P\left( i,j \right) \right)^{2}}{{Nz}^{2}}$ | It measures the variability of gray-level intensity values in the image. It is the normalized version of the GLN formula.  A low value indicates a greater similarity in the intensity values. |
| 89 | Size Zone Non Uniformity (SZN) | $\frac{\sum_{j=1}^{Ns} \left( \sum_{i=1}^{Ng} P\left( i,j \right) \right)^{2}}{Nz}$ | It measures the variability of size zone volumes in the image.   A low value indicates more homogeneity in size zone volumes. |
| 90 | Size Zone Non Uniformity Normalized | $\frac{\sum_{j=1}^{Ns} \left( \sum_{i=1}^{Ng} P\left( i,j \right) \right)^{2}}{{Nz}^{2}}$ | It measures the variability of size zone volumes throughout the image. This is the normalized version of the SZN formula.  A low value indicates more homogeneity among zone size volumes in the image. |
| 91 | Zone Percentage | $ZP=\frac{Nz}{Np}$ | *ZP* measures the coarseness of the texture by taking the ratio of number of zones and number of voxels in the ROI.  Values are in range$\frac{1}{Np}\leq ZP\leq1$.   High values indicate a large portion of the ROI consisting of small zones (indicative of a more fine texture). |
| 92 | Gray Level Variance (GLV) | $GLV=\sum_{i=1}^{Ng} \sum_{j=1}^{Ns} p\left( i,j \right)\left( i-\mu\right)^{2}$ $\mathrm{where}$ $\mu=\sum_{i=1}^{Ng} \sum_{j=1}^{Ns} p\left( i,j \right)i$ | GLV measures the variance in gray level intensities for the zones. |
| 93 | Zone Variance (ZV) | $\sum_{i=1}^{Ng} \sum_{j=1}^{Ns} p\left( i,j \right)\left( j-\mu\right)^{2}$ $where:$ $\mu=\sum_{i=1}^{Ng} \sum_{j=1}^{Ns} p\left( i,j \right)j$ | *ZV* measures the variance in zone size volumes for the zones. |
| 94 | Zone Entropy (ZE) | $-\sum_{i=1}^{Ng} \sum_{j=1}^{Ns} p\left( i,j \right){log}_{2}\left( p\left( i,j \right)+\epsilon\right)$ | ZE measures the uncertainty/ randomness in the distribution of zone sizes and gray levels.  Here, ϵ is an arbitrarily small positive number  $\left( \approx2.2\times{10}^{-16} \right)$.  A higher value indicates more heterogeneneity in the texture patterns. |
| 95 | Low Gray Level Zone Emphasis (LGLZE) | $\frac{\sum_{i=1}^{Ng} \sum_{j=1}^{Ns} \frac{P\left( i,j \right)}{i^{2}}}{Nz}$ | LGLZE measures the distribution of lower gray-level size zones.  A high value indicates a greater proportion of lower gray-level values and size zones in the image. |
| 96 | High Gray Level Zone Emphasis (HGLZE) | $\frac{\sum_{i=1}^{Ng} \sum_{j=1}^{Ns} P\left( i,j \right)i^{2}}{Nz}$ | HGLZE measures the distribution of the higher gray-level values.  A high value indicates a greater proportion of higher gray-level values and size zones in the image. |
| 97 | Small Area Low Gray Level Emphasis (SALGLE) | $\frac{\sum_{i=1}^{Ng} \sum_{j=1}^{Ns} \frac{P\left( i,j \right)}{i^{2}j^{2}}}{Nz}$ | SALGLE measures the proportion in the image of the joint distribution of smaller size zones with lower gray-level values. |
| 98 | Small Area High Gray Level Emphasis (SAHGLE) | $\frac{\sum_{i=1}^{Ng} \sum_{j=1}^{Ns} \frac{P\left( i,j \right)i^{2}}{j^{2}}}{Nz}$ | SAHGLE measures the proportion in the image of the joint distribution of smaller size zones with higher gray-level values. |
| 99 | Large Area Low Gray Level Emphasis (LALGLE) | $\frac{\sum_{i=1}^{Ng} \sum_{j=1}^{Ns} \frac{P\left( i,j \right)j^{2}}{i^{2}}}{Nz}$ | LALGLE measures the proportion in the image of the joint distribution of larger size zones with lower gray-level values. |
| 100 | Large Area High Gray Level Emphasis (LAHGLE) | $\frac{\sum_{i=1}^{Ng} \sum_{j=1}^{Ns} P\left( i,j \right)i^{2}j^{2}}{Nz}$ | LAHGLE measures the proportion in the image of the joint distribution of larger size zones with higher gray-level values. |

#### Neighbouring Gray Tone Difference Matrix (NGTDM) Features

A Neighbouring Gray Tone Difference Matrix quantifies the difference between a gray value and the average gray value of its neighbours within distance *δ*. The sum of absolute differences for gray level *i* is stored in the matrix.

Let:

- $n_{i}$ be the number of voxels in $X_{gl}$ with gray level $i$.
- $N_{v,p}$be the total number of voxels in $X_{gl}$ and equal to $\sum n_{i}$ (i.e. the number of voxels with a valid region; at least 1 neighbor). $N_{v,p}\leq N_{p}$, where $N_{p}$ is the total number of voxels in the ROI.
- $p_{i}$ be the gray level probability and equal to $\frac{}{Nv}$.
- $s_{i}=\left\{ \begin{aligned} \sum_{i=1}^{n_{i}} \left| i-\acute{A}_{i} \right|,\wedge forn_{i}\neq0 \\ 0,forn_{i}=0 \end{aligned} \right.$be the sum of absolute differences for gray level $i$.
- $N_{g}$ be the number of discreet gray levels.
- $N_{g,p}$ be the number of gray levels where $p_{i}\neq0$.

| S. No | Feature Name | Formula | Description |
| --- | --- | --- | --- |
| 101 | Coarseness | $\frac{1}{\sum_{i=1}^{Ng} p_{i}s_{i}}$ | Coarseness is a measure of average difference between the center voxel and its neighborhood. It is an indication of the spatial rate of change.  A high value indicates a low spatial change rate and a locally more uniform texture.   The value of $\sum_{i=1}^{Ng} p_{i}s_{i}$ potentially evaluates to 0 (in case of a completely homogeneous image). If this is the case, an arbitrary value of ${10}^{6}$ is returned. |
| 102 | Contrast | $\left( \frac{1}{N_{g,p}\left( N_{g,p}-1 \right)}\sum_{i=1}^{Ng} \sum_{j=1}^{Ng} p_{i}p_{j}\left( i-j \right)^{2} \right)\left( \frac{1}{N_{v,p}}\sum_{i=1}^{Ng} s_{i} \right)$ $\mathrm{where}p_{i}\neq0$ and $p_{j}\neq0$ | Contrast is a measure of the spatial intensity change, but is also dependent on the overall gray level dynamic range.  Contrast is high when both the dynamic range and the spatial change rate are high. An image with a large range of gray levels, with large changes between voxels and their neighborhood will have a high contrast.  In case of a completely homogeneous image ($N_{g,p}=1$) it would result in a division by 0. In this case, an arbitrary value of 0 is returned. |
| 103 | Busyness | $\frac{\sum_{i=1}^{Ng} p_{i}s_{i}}{\sum_{i=1}^{Ng} \sum_{j=1}^{Ng} \left\vert ip_{i}-jp_{j} \right\vert}$ $\mathrm{where}p_{i}\neq0$ and $p_{j}\neq0$ | Busyness is a measure of the change from a pixel to its neighbor.   A high value for busyness indicates a ‘busy’ image, with rapid changes of intensity between pixels and its neighborhood.   If $N_{g,p}=1$, then $busyness=\frac{0}{0}$. If this is the case, 0 is returned, as it concerns a fully homogeneous region. |
| 104 | Complexity | $\frac{1}{N_{v,p}}\sum_{i=1}^{Ng} \sum_{j=1}^{Ng} \left\vert i-j \right\vert\frac{p_{i}s_{i}+p_{j}s_{j}}{p_{i}+p_{j}}$ $\mathrm{where}p_{i}\neq0$ and $p_{j}\neq0$ | An image is considered complex when there are many primitive components in the image, i.e. if the image is non-uniform and there are many rapid changes in gray level intensity. |
| 105 | Strength | $\frac{\sum_{i=1}^{Ng} \sum_{j=1}^{Ng} \left( p_{i}+p_{j} \right)\left( i-j \right)^{2}}{\sum_{i=1}^{Ng} s_{i}}$ $\mathrm{where}p_{i}\neq0$ and $p_{j}\neq0$ | Strength is a measure of the primitives in an image.  Its value is high when the primitives are easily defined and visible, i.e. an image with slow change in intensity but larger coarse differences in gray level intensities.  If $\sum_{i=1}^{Ng} s_{i}$ evaluates to 0 (in case of a completely homogeneous image) then 0 is returned. |

### Image Filters Applied

The image of the brain before applying any filters is as follows:

The following table contains the filters that are applied on the image:

| S. No | Image Filter Name | Description |
| --- | --- | --- |
| 1 | Additive Gaussian Noise Image Filter | This alters the image with additive Gaussian white noise which can be modelled as:  $I=I_{0}+N$, where I is the observable image, I_0_ is the noise free image and N is a normally distributed random variable of mean and standard deviation. The noise is independent of the pixel intensities. |
| 2 | Binomial Blur Image Filter | This filter performs a separable blur on each dimension of the image.   The binomial blur consists of a nearest neighbor average along each image dimension. The net result after n-iterations approaches convulsion with a Gaussian. |
| 3 | Box Mean Image Filter | Implements a fast rectangular mean filter using the accumulator approach. |
| 4 | Box Sigma Image Filter | Box Sigma Image Filter also implements a fast rectangular sigma filter using the accumulator approach |
| 5 | Curvature Flow Image Filter | This filter implements a curvature driven image de-noising algorithm. Iso-brightness contours in the grayscale input image are viewed as a level set. The level set is then evolved using a curvature-based speed function:  $I_{t}=k\left\vert\nabla I \right\vert$, where *k* is the curvature.  The advantage of this approach is that sharp boundaries are preserved with smoothing occurring only within a region. |
| 6 | Discrete Gaussian Image Filter | Blurs an image by separable convolution with discrete Gaussian kernels. This filter performs Gaussian blurring by separable convolution of an image and a discrete Gaussian operator (kernel). |
| 7 | Laplacian Sharpening Image Filter | This filter sharpens an image using a Laplacian. Laplacian Sharpening highlights regions of rapid intensity change and therefore highlights or enhances the edges. The result is an image that appears more in focus. |
| 8 | Mean Image Filter | It applies an averaging filter to an image.  It computes an image where a given pixel is the mean value of the pixels in a neighborhood about the corresponding input pixel.  Mean filter is a part of the linear filters family. |
| 9 | Median Image Filter | It applies a median filter to an image.  It computes an image where a given pixel is the median value of the pixels in a neighborhood about the corresponding input pixel.  It is used to smooth an image without being biased by outliers or shot noise.  A median filter is a part of the nonlinear filters family. |
| 10 | Normalize Image Filter | Normalize an image by setting its mean to zero and variance to one.  Normalize Image Filter shifts and scales an image so that the pixels in the image have a zero mean and unit variance.   This filter uses *Statistics Image Filter* to compute the mean and variance of the input and then applies *Shift Scale Image Filter* to shift and scale the pixels. |
| 11 | Recursive Gaussian Image Filter | It is the base class for computing IIR convolution with an approximation of a Gaussian kernel and is given by:  $\frac{1}{\sigma\sqrt{2\pi}}exp(-\frac{x^{2}}{2\sigma^{2}})$ $Compared to the Discrete Gaussian Image Filter$ (above), this filter tends to be faster for large kernels, and it can take the derivative of the blurred image in one step. |
| 12 | Shot Noise Image Filter | It alters an image with shot noise. The shot noise follows a Poisson distribution:  $I=N(I_{0})$where $N(I_{0})$ is a Poisson-distributed random variable of mean $I_{0}$.   The noise is thus dependent on the pixel intensities in the image. The Poisson-distributed variable 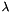 is computed by using the following algorithm: 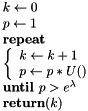 where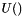 provides a uniformly distributed random variable in the interval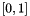.  This algorithm is very inefficient for large values of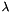. Fortunately, the Poisson distribution can be accurately approximated by a Gaussian distribution of mean and variance 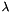 when 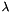 is large enough. Here, this value is considered to be 50. This leads to the faster algorithm:  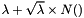  where 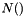 is a normally distributed random variable of mean 0 and variance 1. |
| 13 | Speckle Noise Image Filter | Alter an image with speckle (multiplicative) noise.  The speckle noise follows a gamma distribution of mean 1 and standard deviation provided by the user. The noise is proportional to the pixel intensity.  It can be modeled as:  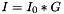  where 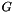 is a gamma distributed random variable of mean 1 and variance proportional to the noise level, *G*:  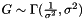 |

### Additional General Information Features

Some additional features were extracted which do not hold significance for the machine learning classifier and they are as follows:

**General Info**

1. Bounding Box": [],
2. Enabled Image Types": {},
3. General Settings": {
   a) "distances": [1],
   b) "additional Info": true,
   c) "force2D": false,
   d) "interpolator": "sitkBSpline",
   e) "voxel Based": false,
   f) "resampled Pixel Spacing": null,
   g) "label": 1,
   h) "normalize Scale": 1,
   i) "normalize": false,
   j) "force 2D dimension": 0,
   k) "remove Outliers": null,
   l) "minimum ROI Size": null,
   m) "minimum ROI Dimensions": 1,
   n) "pre Crop": false,
   o) "resegment Range": null,
   p) "pad Distance": 5},
4. Image Hash
5. Image Spacing
6. Mask Hash
7. Numpy Version
8. PyWavelet Version
9. SimpleITK Version
10. Version
11. Volume Num
12. Voxel Num
